# Severe warming alters biomass allocation and enhances ecosystem respiration in an alpine grassland: evidence from a short-term mesocosm experiment

**DOI:** 10.64898/2026.09.21.753258

**Authors:** Federica D’Alò, Angela Augusti, Carlotta Volterrani, Maurizio Sarti, Alexandru Milcu, Sebastien Devidal, Clement Piel, Leonardo Montagnani, Ilaria Fracasso, Olga Gavrichkova

## Abstract

**Background and aims:** Alpine ecosystems are highly sensitive to climate change, with rising temperatures threatening ecosystem structure and function. This study examined the short-term responses of alpine grassland ecosystems (Mont Blanc, Italy) to projected climate scenarios using an Ecotron mesocosm experiment.

**Methods:** Ecosystem monoliths were exposed to current (~420 ppm CO□) and future CO□ concentrations (~550 and ~800 ppm) under RCP 4.5 and RCP 8.5 scenarios, incorporating changes in temperature, precipitation, humidity, and radiation. CO□ fluxes (*NEP, GPP, R*_*eco*_), soil chemistry, vegetation coverage, above- and belowground biomass, and plant functional traits were assessed throughout the experiment.

**Results:** Alpine grassland ecosystems withstood moderate warming (RCP 4.5), while under severe warming (RCP 8.5), *GPP*, vegetation cover and biomass increased with a shift toward aboveground allocation, although daytime *NEP* remained stable as gains were offset by higher *R*_*eco*_, with no detectable water and nutrients limitations. On a diel basis, higher GPP was offset by enhanced *R*_*eco*_, indicating a potential towards net C loss. *Salix herbacea* exhibited functional trait adjustments—higher LMA and N_a_, and elevated δ^1^□N—reflecting greater leaf longevity, water-use efficiency, and altered nitrogen metabolism, suggesting potential community reorganization.

**Conclusion:** Over a single growing season, alpine ecosystems appeared relatively stable under moderate warming. However, under the more severe RCP 8.5 scenario, biomass allocation shifted toward greater aboveground and reduced belowground investment, leading to altered C fluxes that, if sustained over the long-term, could drive the ecosystem C balance from a sink to a source. Responses were plant species-specific, indicating uneven acclimation within the community.

## Introduction

Alpine ecosystems are among the most sensitive and vulnerable regions to climate change with temperatures rising about twice as fast as the global average (Kotlarski et al. 2023). Characterized by low temperatures and pronounced seasonality, these ecosystems support unique biodiversity adapted to harsh conditions, where plant growth is primarily constrained by temperature (Ma et al. 2010). Mean annual temperatures (MAT) in the Alps have risen by about +1.8°C over the past 150 years, with accelerated warming since the 1980s, and are projected to rise by an additional 2-4°C by the end of the century (IPCC 2021).

Mountain grasslands, which cover 20–25% of terrestrial ecosystems are among the most climate-sensitive biomes and represent important global soil carbon (C) reservoirs due to the accumulation of organic matter under low-temperature conditions (Török et al. 2018, Hirota et al., 2009). Ongoing climate changes alter ecosystem functioning, thereby influencing C, water and nutrient cycles, as well as plant and microbial community composition. In particular, changes in Gross Primary Productivity (*GPP*) and Ecosystem Respiration (*R*_*eco*_) jointly determine Net Ecosystem Productivity (*NEP*), thereby regulating whether ecosystems act as carbon sinks or sources (Rogora et al. 2018; Dong et al. 2022). However, predicting alpine ecosystem responses to climate change remains challenging due to the complex interplay of multiple climatic drivers and the biological interactions they trigger.

Temperature manipulation experiments indicate that ecosystem responses to climate change in cold environments differ from those observed in temperate and arid regions. In these ecosystems, temperatures are typically below the metabolic optimum and warming-induced drought is partly buffered by snow-derived soil moisture (Volk et al., 2021). In particular, under moderate warming, plant growth and activity are strongly stimulated. However, under more severe warming, this positive response is attenuated due to increasing water limitation. Despite this limitation, overall plant productivity during the growing season remains positive (Volk et al., 2021; Volk et al., 2022). Additionally, the beneficial effects of warming on C uptake may be constrained at extreme temperatures by mismatches between the temperature optima of photosynthesis and plant respiration (Gavrichkova et al., 2022). Warming-induced increases in plant productivity are often associated with enhanced belowground C allocation, which can stimulate root respiration, both by increasing substrate supply to existing roots and by promoting new root biomass formation (Bahn et al., 2008). At the same time, warming enhances heterotrophic respiration, driven by microbial decomposition of labile soil C and root exudates. The stimulation of microbial activity also enhances nitrogen (N) mineralization, leading to increased soil N availability that supports plant growth (Gavrichkova et al., 2017).

However, this effect is generally reported to be short-lived, as microbial responses to sustained warming tend to attenuate over time. This apparent adaptation is driven by a combination of physiological adjustments and shifts in microbial community composition. It is also associated with the progressive depletion of readily decomposable substrate and warming-induced reductions in soil moisture. Together, these factors constrain microbial processes and ultimately slow respiration rates (Fei et al., 2015; Walker et al., 2018; Tiwari et al., 2021).

Plant species also respond differently to warming, depending on individual physiological plasticity and tolerance to increasing temperature and change in hydrological cycle (Cannone et al. 2016). Species-specific responses are often accompanied by changes in leaf functional traits, including adjustments in structural properties and leaf nutritional status, reflecting their acclimation strategies. Nevertheless, the ecophysiological response to climate change as expected by function traits adjustment, remain poorly documented across different plant life forms (Yu et al. 2022).

The evidences discussed above are largely derived from experimental warming studies in which temperature is manipulated in isolation, which limits their ability to represent and predict ecosystem responses to the concurrent and interacting changes characteristic of real-world climate change. Until recently, simulating realistic future conditions in climate change experiments was not feasible (Korell et al. 2019), as such experiments necessitate precise control over environmental variables to replicate both current and future climate conditions. Advanced controlled environment facilities for ecosystem research (Ecotrons) fulfill these needs by offering systems that can simultaneously manipulate and measure multiple parameters (Roy et al. 2016; Roy et al. 2022).

In this context, this study employed an Ecotron experimental platform to assess how alpine grassland monoliths respond to future climate scenarios, linking CO_2_ fluxes with changes in vegetation coverage and species-specific leaf functional traits. Measurements targeted representative daytime ecosystem gas exchange under sustained photosynthetic activity, with *NEP* providing an estimate of the potential daytime C-sink strength. To this end, ecosystem samples/monoliths extracted from an alpine site in Mont Blanc area, Italy, were exposed to short-term controlled climate simulations, representing current (~420 ppm CO□) and future projected atmospheric CO□ concentrations (~550 ppm and ~800 ppm) aligned with mid-century projections under the representative concentration pathway (RCP) 4.5 and RCP 8.5 emission scenarios. These simulations took into account respective changes in key environmental variables, including air temperature, precipitation, air relative humidity, and short-wave radiation.

We hypothesized that: *i*) Under moderate climate change scenarios (RCP 4.5), enhanced temperatures and CO_2_ rise will stimulate photosynthesis (GPP), followed by enhanced plant growth expressed here as aboveground biomass and vegetation coverage. These changes are expected to be accompanied by an increase in *R*_*eco*_, resulting in daytime *NEP* that remains neutral to positive throughout the growing season compared to control conditions. These responses will coincide with a gradual increase in nutrients availability in the soil due to accelerated microbial activity; *ii*) Under severe warming (RCP 8.5), particularly when sustained by limited water availability, plant growth and GPP are expected to show only modest increase. *R*_*eco*_ may be constrained by substrate and moisture limitations, leading to a reduced nutrient pool associated with decreased microbial activity compared to control conditions. This will result in higher daytime *NEP* due to enhanced C assimilation and reduced *R*_*eco*_; iii) Plant responses to climate scenarios would be species-specific, reflected in interspecific differences in leaf functional traits and growth dynamic.

## Material and Methods

### Description of the sampling area

The study area is located near the Col de la Seigne in Val Veny, on the southwestern slope of the Mont Blanc Massif, at an altitude of approximately 2,500□m (45°45’03.6”N, 6°48’25.2”E) (Figure 1). The site faces east and has a slope of roughly 20°. Mean annual temperatures in the last ten years ranged between 0.7□°C and 2.6□°C, with winter minima occasionally dropping below −15□°C and summer maxima rarely exceeding 20°C (data of Centro Funzionale RAVDA https://presidi2.regione.vda.it/l accessed 2025 for the closest stations at similar elevation). Vegetation is dominated by *Potentilla aurea, Salix herbacea, Scorzoneroides montana, Poa alpina*, and *Carex foetida*. The area is characterized by the prevalence of shallow Leptosols and locally occurring silty deposits (DataBasin mapping platform, Conservation Biology Institute, https://databasin.org, accessed 2025). The soil was previously identified as sandy loam, consisting of approximately 71% sand, 24% silt and 5.4 clay (Fracasso et al., 2025). To further characterize site-specific soil physical macroscopic properties, fifteen additional soil samples were collected from the 0–15 cm soil layer (unpublished data). Briefly, soil samples were oven-dried at 60 °C and sieved through a 2 mm mesh to separate fine soil, stones, and belowground organic components. Each fraction was then weighed separately to quantify its contribution to total sample dry mass. In the upper 10 cm of soil, the average stone fraction was 31.7%, the fine-soil fraction 65.9%, and belowground plant biomass (roots and rhizomes) 2.4%. At 10–15 cm depth, stones became dominant, while belowground plant biomass decreased to 0.4%, with 51.9% stones and 47.7% fine soil.

**Fig 1.**
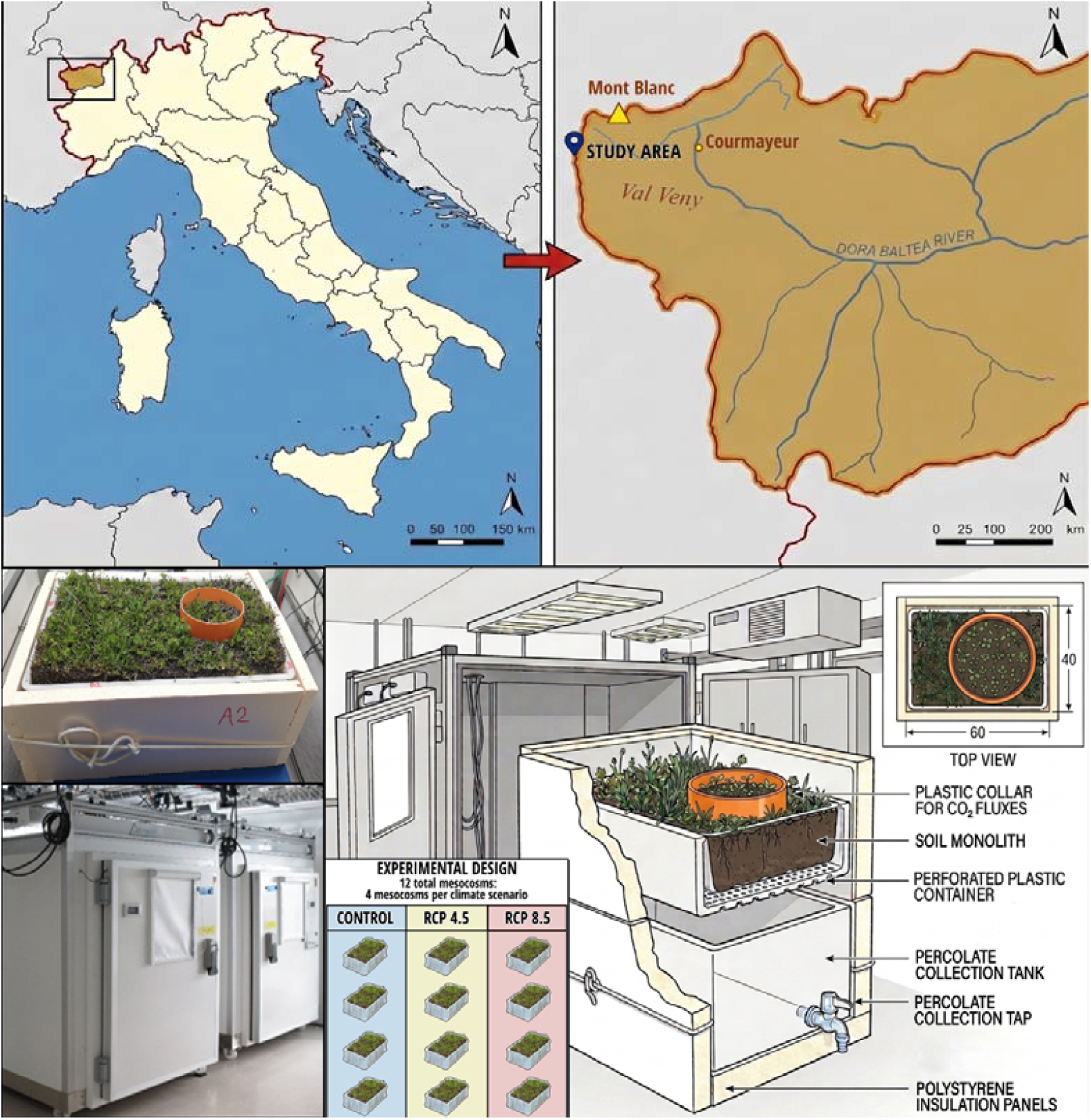
Sampling area map (top) and schematic representation of the mesocosm experiment setup (bottom). The figure was partially generated using Gemini AI and subsequently edited.

### Mesocosm experimental setup

Alpine ecosystem monoliths (60 cm length × 40 cm width × 12 cm height) were collected from the high-altitude alpine site at the beginning of the vegetation season, immediately after snowmelt. The monoliths were carefully extracted from the topsoil using spades and shovels, ensuring minimal disturbance to the vegetation. Depth of the monoliths was chosen experimentally, to ensure that 80% of root biomass was included. Each monolith, representative of the wider sampling area, included a balanced composition of the dominant plant types, including *Potentilla aurea, Salix herbacea, Scorzoneroides montana, Poa alpina* and *Carex foetida*. The monoliths were carefully placed into perforated plastic containers directly in the field for transport and experimental use, ensuring the maintenance of air and water flow. The perforations allowed the passage of air and water, preventing excessive moisture accumulation or stagnation, which could have otherwise altered the microbiological or chemical composition of the samples. A total of 12 monoliths were transported under controlled temperature conditions to the Mesocosms Experimental Platform of the Montpellier European Ecotron (CNRS, France), where they were randomly distributed across the 12 controlled environment chambers/phytotrons, with four chamber replicates for each of the three climate scenarios (Figure 1). The climate chambers had a total volume of 2□m^3^ and a working surface area of 1□m^2^. The monoliths, placed in perforated containers, were then positioned on top of a water collecting tanks, from which the water was periodically emptied to avoid water accumulation underneath. The entire setup was insulated with polystyrene panels lining the container to minimize heat loss during chamber opening for maintenance and measurement activities in the Ecotron. The climate scenarios included a Control, reflecting current climate conditions (~420 ppm CO_2_) and two future projections RCP 4.5 and RCP 8.5 (~550 ppm CO_2_ and ~800 ppm CO_2_, respectively), simulating conditions forecasted for mid-century (2051-2070). Within the chambers, climate variables— air temperature, precipitation, air relative humidity, photosynthetically active radiation (PAR; 0–1200 µmol m□^2^ s□^1^), and CO_2_ concentrations—were systematically manipulated with a 30 sec time step to replicate the projected scenarios. A two-week acclimation period was allowed before starting the measurements to minimize the effects of transport-induced stress on the monoliths. The total experimental duration, including the acclimation phase, was 57 days.

### Climate scenarios simulation within climate chambers

Current and future climate scenarios for the chamber experiments were generated using hourly data from the original Alpine monolith location, provided by the Euro-Mediterranean Center on Climate Change (CMCC) (Raffa al. 2021; Adinolfi et al. 2023). Two high-resolution datasets (2.2 km) over Italy were used: the ERA5 reanalysis dataset (“ERA5 downscaling 2.2 km over Italy”, 1991–2020) and a climate projection dataset based on RCP4.5 and RCP8.5 scenarios (“Climate Projections RCP4.5 and RCP8.5 downscaled at 2.2 km over Italy”, 2001–2070).*The delta change (ΔC) approach (Hay et al., 2000) was used to simulate three climate scenarios for the experiment. The Control scenario, representing current climate conditions, was defined as the mean climate of the ten warmest years within the ERA5 dataset (1991–2020). Two future scenarios, Future RCP4.5 and Future RCP8.5, represent mid-century climate conditions and were calculated as 20-year averages for the period 2051–2070.

Future climate scenarios were obtained by adding ΔC to the Control scenario, according to the following equations:

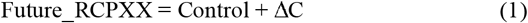

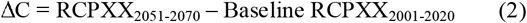

*where XX indicates either 4*.*5 or 8*.*5. Here, RCPXX*□□□□□□□□□ *is the mean projection for 2051-2070 and* Baseline *RCPXX*□□□□□□□□□ *represents the 20-year reference period (2001-2020), obtained by combining historical simulations (2001–2005) with the corresponding early RCPXX data (2006–2020) within the climate projections dataset*.

The simulations included air temperature, precipitation, relative humidity, solar radiation and CO_2_ concentrations. All parameters were simulated at an hourly time step, except for precipitation, which was simulated on a weekly basis by applying the prescribed amount of water through irrigation. Precipitation was manually supplied by irrigating the mesocosms with water purified by reverse osmosis and UV disinfection. CO□ concentrations were fixed at 420, 550, and 800 ppm for the Control, RCP4.5, and RCP8.5 scenarios, respectively.

### Soil physicochemical characterization

Soil was collected at the beginning of the experiment and subsequently every 20 days using a sterilized spatula from at least three points within each mesocosm, up to a depth of 10□cm. Samples from the different points were mixed to obtain a composite sample and transferred into sterile Falcon tubes, which were immediately frozen and stored at −20□°C until analysis. To analyze the chemical composition of soil, samples were air-dried until they reached a constant weight and then sieved through a 2 mm mesh. Soil physicochemical parameters, including pH, total carbon (TC), total nitrogen (TN), nitrate (NO□□), ammonium (NH□□), available phosphate (PO□^3^□), and total phosphorus (TP), were determined as described in the Supplementary Material.

### Flux measurements and calculations

Flux measurements were conducted from 30 June 2023 (Day Of Year, DOY 181) to 7 August 2023 (DOY 219). CO_2_ fluxes were measured twice a week, starting from late morning till early afternoon. Fluxes measured during this period are commonly considered as representative of daytime gas exchange under sustained photosynthetic activity and avoid transient effects associated with dawn and dusk. The optimum time window for measurements was confirmed based on a preliminary diurnal assessment during which fluxes were monitored from nighttime to late afternoon (data not shown). An EGM-5 Portable CO_2_ Gas Analyzer (PP Systems, USA) equipped with a CPY-5 Transparent Canopy Assimilation Chamber with a diameter of 14.6 cm was used to measure the CO_2_ fluxes. Plastic collars, inserted approximately 2 cm into the soil, were installed on each monolith at the start of the experiment upon their arrival at the Ecotron, allowing repeated measurements to be taken at the same location. The chamber was equipped with sensors for the measurement of air temperature and PAR. CO_2_ concentrations were measured at ambient light to estimate the *NEP* of the monoliths and in the dark by shading the chamber to measure *R*_*eco*_. To minimize short-term physiological artifacts associated with the light–dark transition, *R*_*eco*_ was measured after a short period of darkening, once the CO□ flux in the dark reached steady-state conditions. The duration of this stabilization period was determined empirically during preliminary measurements. We used the biological convention in which a positive value of *NEP* represents CO□ uptake (CO□ sink) into the ecosystem, whereas a negative value indicates CO□ release to the atmosphere (CO□ source). *R*_*eco*_ is considered positive, meaning that higher *R*_*eco*_ values correspond to greater CO□ emissions. Gross primary productivity (*GPP*), which represents total photosynthetic C uptake, was then calculated as the sum of *NEP* and *R*_*eco*_ (*GPP* = *NEP* + *R*_*eco*_). Because respiration measured after darkening may differ from respiration occurring under illuminated conditions due to the *Kok effect* or the light inhibition of leaf respiration (Heskel et al., 2013), we acknowledge that *GPP* values derived from such partitioning approach could be overestimated. Nevertheless, this method remains appropriate for comparative analyses across treatments and is commonly adopted in chamber-based studies (Järveoja et al., 2018; Wang et al. 2021), provided that the same measurement protocol is consistently applied.

### Vegetation coverage

Vegetation coverage in the collars, where C fluxes were measured, was assessed using ImageJ software (Schneider et al. 2015). Weekly images of the vegetated surfaces were captured with a high-resolution Nikon Z6II (Japan) digital camera at a fixed distance of 50 cm, under consistent lighting conditions to minimize variability. Images were processed in ImageJ to calculate the proportion of vegetated area by distinguishing vegetation from non-vegetated surface. Vegetation coverage was expressed as the area of vegetation (cm^2^) relative to the total collar area (167 cm^2^), providing a quantitative estimate of vegetation dynamics over time.

### Above- and belowground biomass components

The above- and belowground plant biomass measurements were conducted at the conclusion of the experiment and, involved destructive analyses. Aboveground biomass, litter mass and soil within the collars were harvested for laboratory analysis of both aboveground and belowground components. Plants were separated by dominant species, and a set of plant functional traits was quantified to characterize species-level ecological strategies and resource-use patterns (Wright et al., 2004). Specifically, functional traits included morphological (projected leaf area, species coverage, leaf mass per area) and biogeochemical traits (leaf N concentration expressed on a mass and area basis, and isotopic composition, δ^1^□N). The projected leaf area for each species was measured using ImageJ software version 1.54g. The coverage of each plant species was calculated relative to the collar area as a percentage. The samples were then dried in an oven at 60°C for 72 hours and weighed. Then, samples were ground into a homogeneous powder using a ball mill (MM 400, Retsch) and N concentrations and isotopic composition (δ ^15^N) of leaves from different vegetation species were analyzed (Isoprime, GV Instruments coupled with NA 1500 CNS Analyzer, Carlo Erba). Then, leaf mass per area (LMA) was calculated by dividing the dry leaf mass (grams of dry weight) by the projected leaf area (m^2^). N per leaf mass (N_m_) was calculated as the percentage of N content per gram of dry leaf mass, while N per leaf area (*N*_*a*_) was calculated as (N_m_/100)*LMA. The total LMA was measured by dividing the sum of dry leaves mass by the projected leaves area (m^2^).

Soil samples were sieved to 2 mm, roots were carefully extracted, washed from soil dried, and weighed. The belowground to aboveground biomass ratio was calculated by dividing root biomass by the sum of aboveground and litter biomass.

Soil basal respiration was measured following the method described by Creamer et al. (2014), with modifications optimized and validated in our laboratory. Briefly, 1 g of sieved soil was pre-incubated under standardized conditions (25°C and 60% soil water holding capacity) for 5 days. Once the respiration activity reached the steady state, vials with soil were closed and flushed with the CO_2_ free air for 5 min. After 3 hours of incubation the CO□ formed in the headspace of the vials was analyzed using a Multiflow system (Gilson) coupled to an isotope ratio mass spectrometer (Isoprime, GV Instruments). The concentration of the CO_2_ in the vials was calculated against the calibration curve, constructed using the mixtures of air with different CO_2_ concentration.

### Statistical analysis

Linear mixed-effects models were fitted using the *nlme* package (Pinheiro et al. 2017) to assess the effects of climate scenario and sampling date on temporal dynamics of *NEP, GPP*, and *R*_*eco*_. The model included fixed effects of climate scenario, sampling date (as a factor), and their interaction. Random intercepts accounted for the nested structure of monoliths (pots) within sites (*random = ~1* | *site/potID*). To address temporal autocorrelation due to repeated measurements on the same pot, a first-order autoregressive correlation structure (*corAR1*) was included. Additionally, heterogeneity of variance across sampling dates was modeled using a variance structure allowing different variances per date (*varIdent* function). Model selection was based on Akaike Information Criterion (AIC). Post-hoc comparisons of estimated marginal means (*emmeans*, Lenth et al. 2018) were performed to explore potential differences among scenarios at individual time points.

The same model was applied to analyze the differences in vegetation coverage and soil physicochemical parameters, incorporating pre-treatment values as a covariate to account for initial variations that may arise from sampling differences.

To identify the environmental parameters most strongly correlated with C fluxes, multiple regression models including all potential predictors were fitted using the *nlme* package in R. Random effects accounted for repeated measurements over time within each site and pot (random = ~ date | site/potID), and temporal autocorrelation was modeled with an *corAR(1)* structure. A model-averaging approach was applied (MuMIn package, Bartoń 2025) considering only models with ΔAIC ≤ 4, and the best predictors were selected based on the highest AIC weight.

Statistical tests for daytime *NEP*, daytime *GPP*, daytime and nighttime *R*_*eco*_ and biomass components were performed using one-way analysis of variance (ANOVA) followed by a post hoc pairwise multiple comparison procedure (Tukey’s HSD). Shapiro-Wilk tests were conducted to assess the normality of the data. When necessary, data transformations were applied to achieve normality.

Statistical analyses were performed with R version 4.2.2 (R Core Team, 2022).

### Climate variables under different climate scenarios

Compared with the Control scenario, mean air and soil temperatures were increased 2 and 3°C, respectively, under RCP 4.5, and by 4 and 5°C, respectively, under RCP 8.5 scenario (Figure 2). Mean relative humidity was 71% under Control, 68% under RCP 4.5, and 63% under RCP 8.5 (Figure 2). As the simulations predicted, the RCP 4.5 and RCP 8.5 treatments incorporated 11% and 28% reductions in average simulated precipitation compared to the Control, respectively. Soil water content did not differ among scenarios in terms of mean values, but showed a general decreasing trend over time across all scenarios (Figure 2). The diurnal temperature range (DTR), defined as the difference between daily maximum and minimum air temperatures and reflecting within-day temperature variability, differed slightly among treatments. Over the entire measurement period, mean DTR values were 7.27□±□1.15□°C for Control, 8.11□±□1.21□°C for RCP 4.5, and 9.02□±□1.32□°C for RCP 8.5.

**Fig 2.**
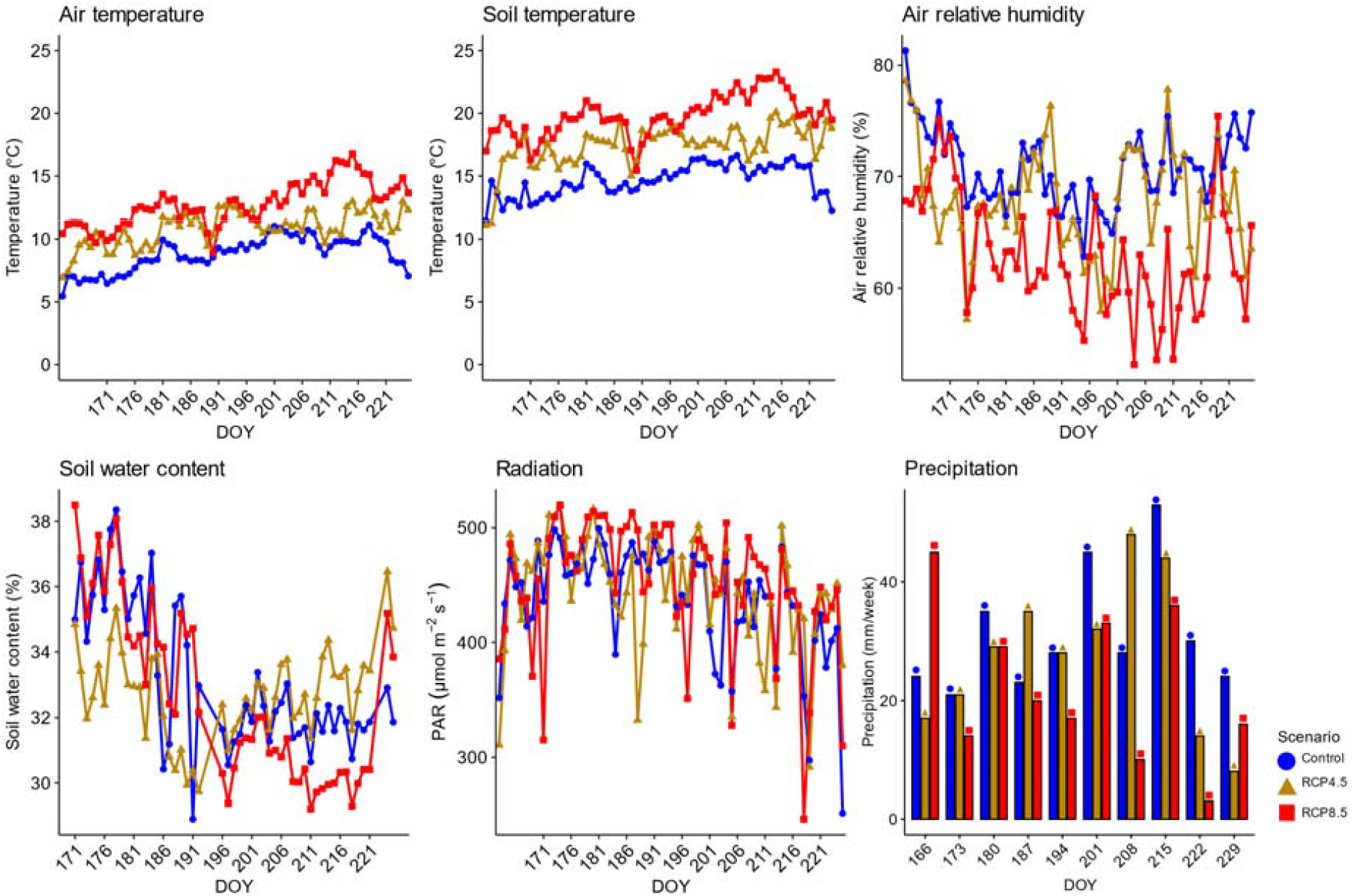
Daily mean values of environmental parameters recorded in the mesocosms for all scenarios, reflecting the conditions imposed by the three climate scenarios. Shown are averages of air temperature, soil temperature, relative humidity, soil water content, and radiation, along with weekly averages of precipitation derived from manual irrigation. DOY = Day Of Year

## Results

### Soil physicochemical characterization

Although some temporal variation was observed, the climate scenarios did not significantly affect soil physicochemical parameters overall. Only PO□^3^□ showed a significant effect of the climate scenarios, likely reflecting initial differences among monoliths and pre-existing heterogeneity among treatments prior to the experiment. Trends in other parameters were generally similar across the Control, RCP 4.5, and RCP 8.5 treatments throughout the experiment (Supplementary Figure 1, Table 1S).

### Impact of climate scenarios on *R*_*eco*_, *GPP* and *NEP*

Figure 3 illustrates the effects of different climate scenarios on key ecosystem processes, with clear differences in *R*_*eco*_ (Table 2S, 3S). *R*_*eco*_ was the highest under the RCP 8.5 scenario, followed by RCP 4.5 and the Control. Accordingly, both climate scenarios (p = 0.006) and their interaction with time (p < 0.001) significantly affected *R*_*eco*_. Soil temperature was positively correlated with *R*_*eco*_, with an increase of 0.181 µmol m□^2^ s□^1^ per 1°C increase of soil temperature.

**Fig 3.**
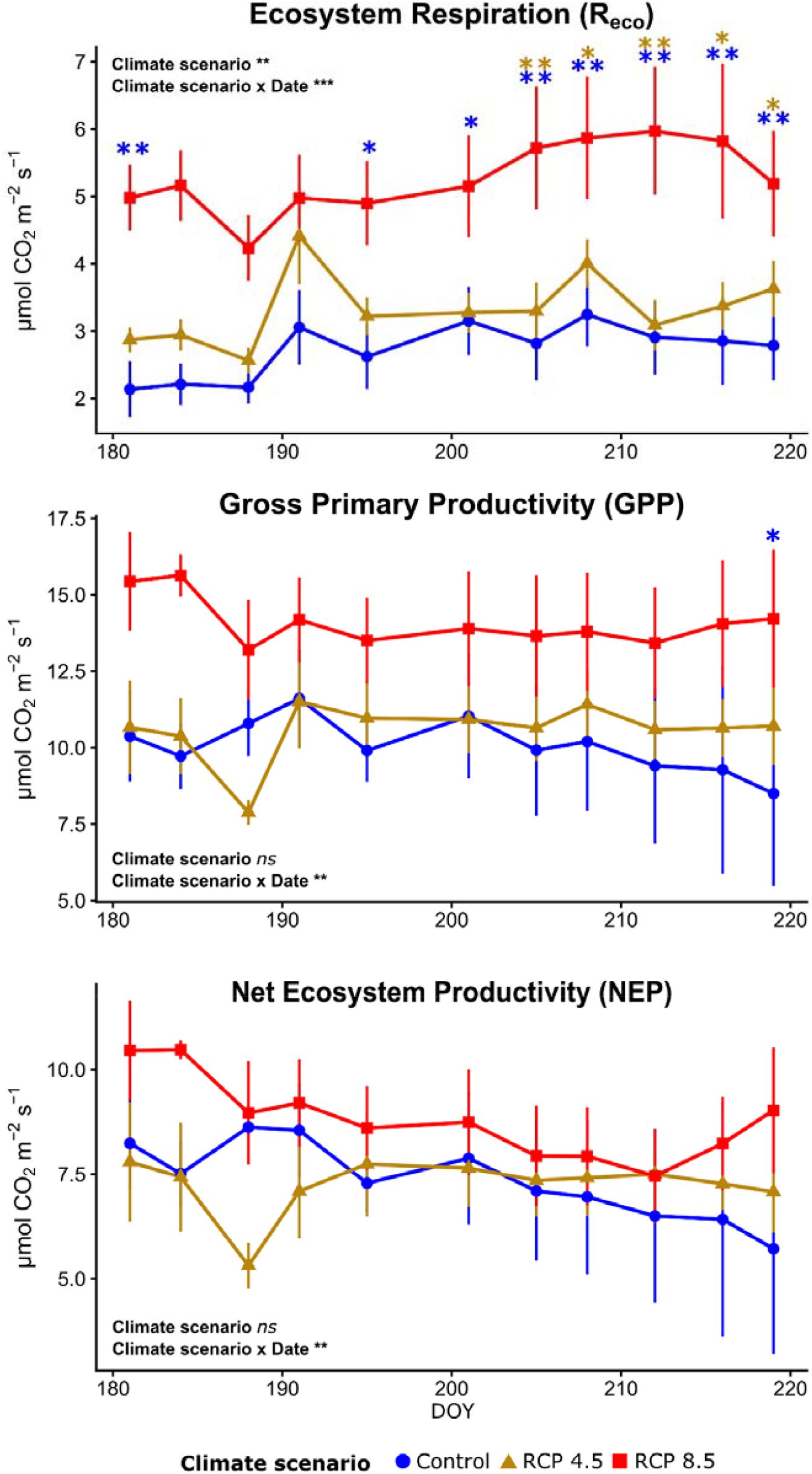
Means ± standard error (n=4) of Ecosystem Respiration, Gross Primary Productivity and Net Ecosystem Productivity under different climate scenarios. Asterisks indicate significant differences based on *emmeans* post-hoc comparisons, with the color showing which scenario the difference refers to. Results from a nonlinear mixed-effects model analyzing carbon fluxes as a function of climate scenario and sampling date are also shown. Significance levels are indicated as *** *p* < 0.001, \*\**p* < 0.01, \**p* < 0.05. DOY = Day Of Year

*GPP* (Table 2S, 3S) showed no significant main effect of climate scenario (p = 0.199). However, a significant interaction between climate scenario and time (*p* = 0.002) indicates that the effect of the scenario varies over time. *GPP* was generally highest under RCP 8.5, although significant differences among scenarios were only observed at the end of the experiment. Air temperature was positively correlated with *GPP*, with an increase of 0.15 µmol m□^2^ s□^1^ per 1 □C increase of air temperature.

*NEP* (Table 2S, 3S) did not differ significantly among climate scenarios, although a significant interaction between climate scenario and time (*p* = 0.006) was detected, suggesting that the effect of climate scenarios on *NEP* changes over time. Air temperature was negatively correlated with *NEP*, with an estimated decrease of 0.027 µmol m□^2^ s□^1^ per 1 □C increase of air temperature.

**Table 1.** Daytime *NEP* and *GPP*, daytime and nighttime *R*_*eco*_ under climate scenarios.

|  | Control | RCP 4.5 | RCP 8.5 |
| --- | --- | --- | --- |
| Daytime $NEP$ | $7.04 \pm 0.46$ **a** | $7.06 \pm 0.52$ **a** | $7.73 \pm 0.47$ **a** |
| Daytime $GPP$ | $9.96 \pm 0.67$ **a** | $10.59 \pm 0.56$ **a** | $12.86 \pm 0.67$ **b** |
| Daytime $R_{eco}$ | $2.92 \pm 0.30$ **a** | $3.53 \pm 0.27$ **a** | $5.14 \pm 0.39$ **b** |
| Nighttime $R_{eco}$ | $2.06 \pm 0.35$ **a** | $2.70 \pm 0.24$ **ab** | $4.04 \pm 0.65$ **b** |

Values are expressed as means ± standard error. The effect of climate scenario on fluxes was assessed using a one-way analysis of variance (ANOVA), followed by Tukey's post hoc test. Different letters represent statistically significant differences at the 0.05 significance level. $NEP$ = Net Ecosystem Productivity; $GPP$ = Gross Primary Productivity; $R_{eco}$ = Ecosystem respiration.

Additionally, daily measurements, summarized in Table 1, were conducted from nighttime through the end of the day. For daytime *NEP*, which reflects the net C exchange during daylight hours, similar values were observed across all scenarios. Indeed, daytime *GPP* and *R*_*eco*_ followed similar trends, significantly higher values under RCP 8.5 compared to both the Control and RCP 4.5. However, nighttime fluxes, represented by *R*_*eco*_ and reflecting the net C exchange during the night, showed significant differences among scenarios. Notably, the RCP 8.5 scenario exhibited more positive values, indicating increased C release at night under these more extreme climate conditions.

### Vegetation coverage and above- and belowground biomass

The vegetation coverage tended to increase under RCP 8.5 (Figure 4, Table 2S). However, the main effect of the climate scenario on vegetated area was not statistically significant, nor was the interaction between the climate scenario and time. While no significant difference was observed in vegetation coverage between RCP 4.5 and the Control, the effect of RCP 8.5 compared to the Control was marginally significant (*p* = 0.085). Additionally, the variable “pre-treatment vegetation coverage” was not statistically significant, indicating that initial differences among plots did not influence vegetation coverage during the experiment.

**Fig 4.**
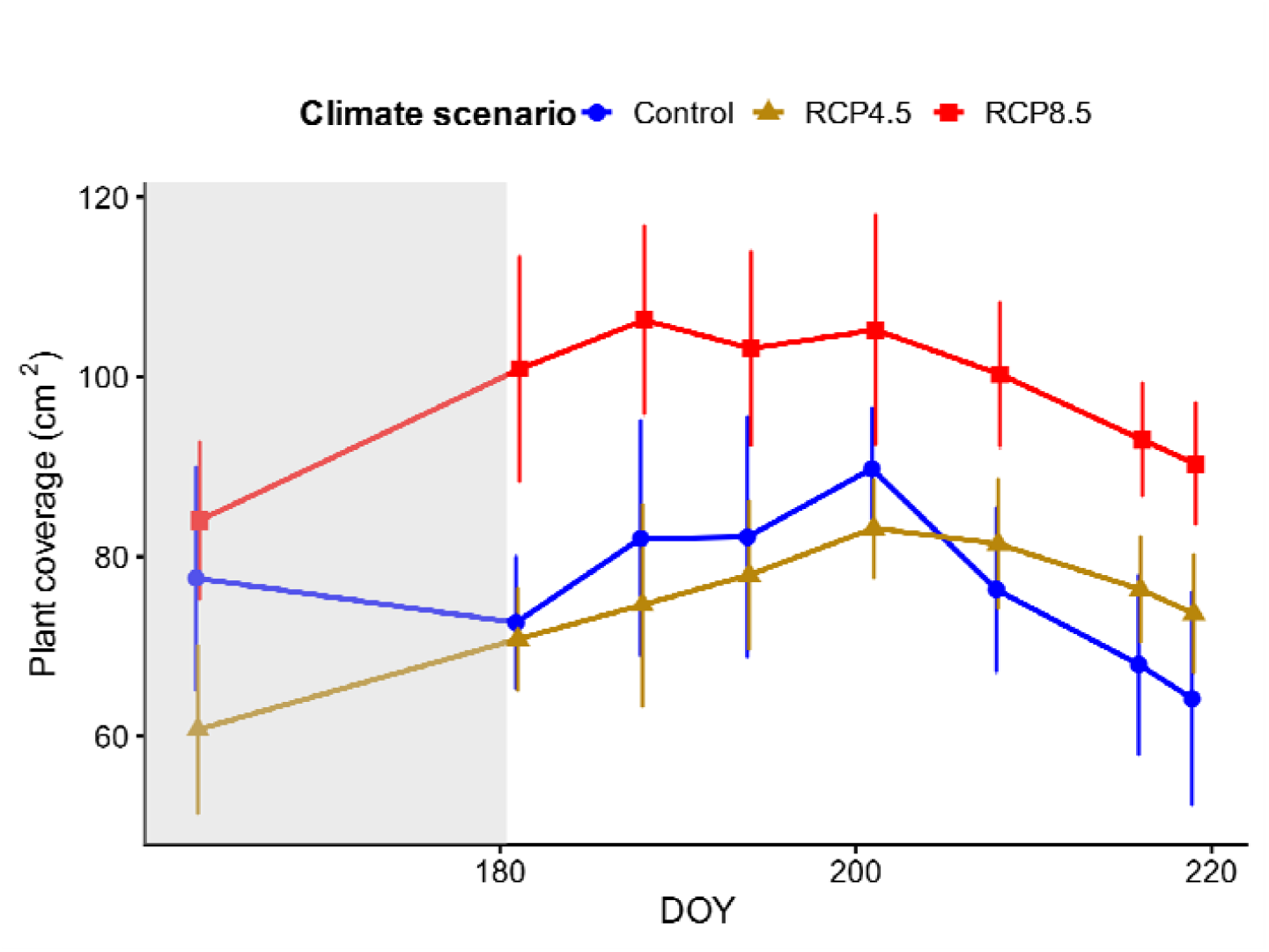
Mean ± standard error of vegetation coverage changes (cm^2^) over time under three climate scenarios. The gray section represents the two-week period prior to the start of measurements. DOY = Day Of Year

**Table 2.** Plant biomass components, LMA and soil basal respiration across climate scenarios.

|  | Control | RCP 4.5 | RCP 8.5 | p-value |
| --- | --- | --- | --- | --- |
| **Aboveground biomass (g DW)** | 3.25±1.89 | 3.61±1.65 | 5.47±1.66 | 0.837 |
| **Litter biomass (g DW)** | 1.30±0.51 | 1.43±0.50 | 1.02±0.33 | 0.806 |
| **Roots (g DW)** | 11.25±2.59 | 13.43±4.04 | 8.08±1.67 | 0.443 |
| **Belowground to aboveground biomass ratio** | 2.85±0.62 | 2.86±0.73 | 1.45±0.28 | 0.084 |
| **LMA (g DW m<sup>-2</sup>)** | 128.62±46.73 | 92.56±13.48 | 86.09±8.09 | 0.944 |
| **Soil basal respiration (µg CO<sub>2</sub>-C g<sup>-1</sup> h<sup>-1</sup>)** | 2.13±1.23 | 2.39±0.35 | 2.57±0.62 | 0.819 |

Values are expressed as means ± standard error (n=4). The effect of climate scenario on the dependent variable was tested using a one-way analysis of variance (ANOVA, p-value). LMA = Leaf Mass Area; g DW = Gram of dry weight

Aboveground biomass, litter biomass, and root biomass were not significantly different between scenarios (Table 2). However, the below/aboveground biomass showed a marginally significant difference (*p* = 0.084) between the scenarios. The LMA and basal respiration did not differ significantly between the scenarios.

Figure 5 presents boxplots illustrating variations in leaf traits across species and climate scenarios at the end of experimental period. Species exhibit distinct patterns in δ15N, with no significant effects of climate scenarios observed within any species. While N_m_ did not differ significantly across scenarios, N_a_ showed significant differences in *Salix herbacea*. Specifically, the RCP 8.5 scenario exhibited higher N_a_ and LMA values compared to the Control, with the RCP 4.5 scenario yielding intermediate values.

**Fig 5.**
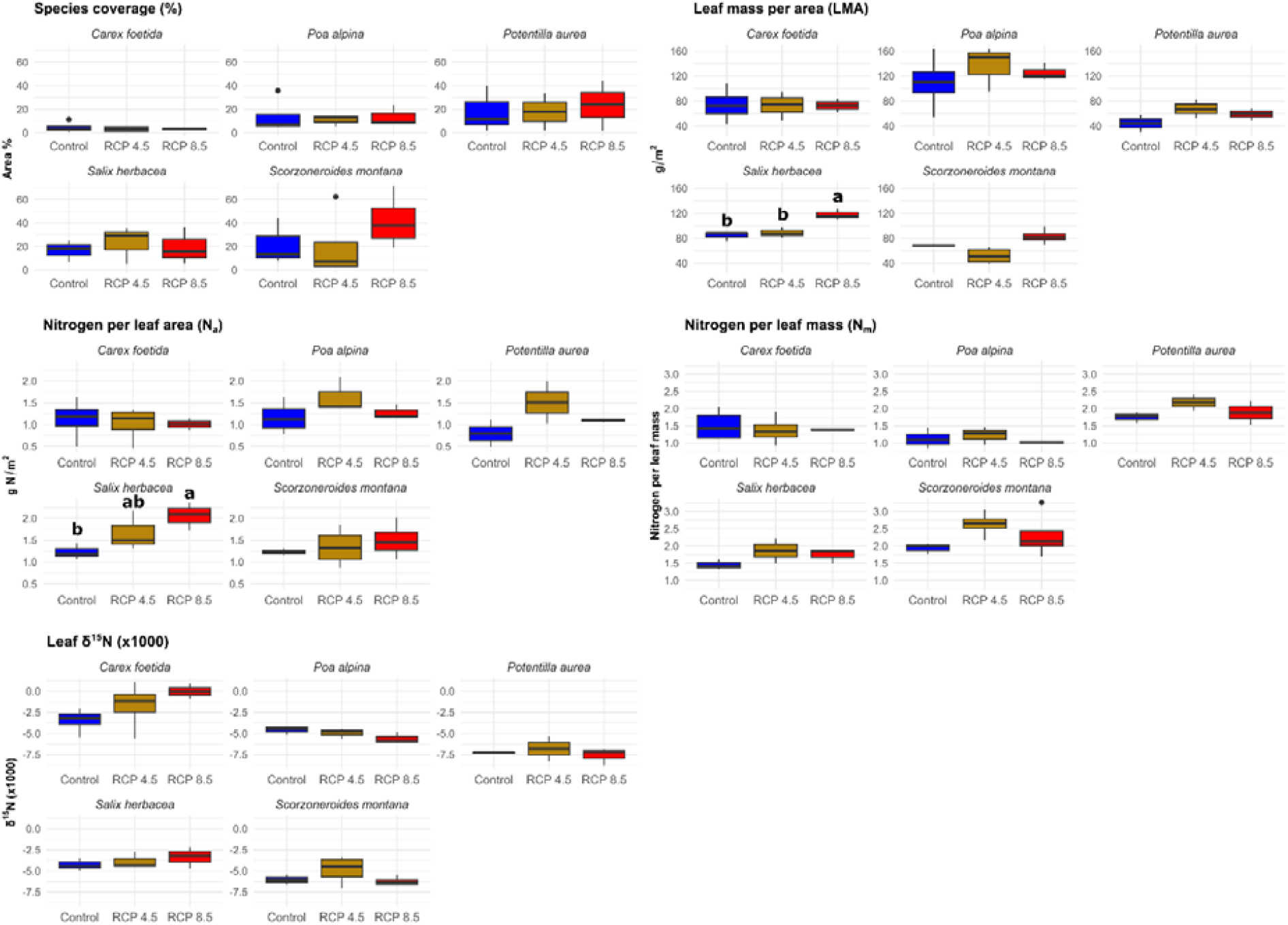
Species-specific plant functional traits different climate scenarios. The species analyzed were *Carex foetida, Poa alpina, Potentilla aurea, Salix herbacea*, and *Scorzoneirides montana*. Letters are reported only for significant differences in one-way ANOVA post hoc Tukey’s HSD test (*p* < 0.05) for Control, RCP 4.5 and RCP 8.5

## Discussion

Alpine grasslands are important C sinks, and understanding how the C balance as a measure of ecosystem resilience will respond to climate change is essential for predicting future ecosystem functioning (Bahn et al., 2006). This study represents one of the few investigations exploring the effects of projected climate change scenarios on alpine grassland ecosystems, using a mesocosm experiment in an Ecotron facility. Such controlled systems bridge theoretical models and real-world complexity, offering a powerful framework for a better understanding of ecosystem processes including nutrient cycling, species interactions, and climate impacts (Luiselli and Pacini, 2025).

### Ecosystem C balance under climate scenarios

Regional climate projections consistently indicate strong and robust changes in the climate of the European Alps by the end of the century, with the magnitude depending on the emission scenario (Gobiet and Kotlarski et al. 2020; Kotlarski et al. 2023). In particular, RCP 8.5 is associated with stronger warming and drier conditions compared to RCP 4.5 and the Control, including reduced relative humidity, summer precipitation, and snow cover. Under RCP 4.5, mean annual temperature is projected to increase by approximately +2 to +4 °C, while under RCP 8.5 warming may reach up to +6 °C, particularly in summer and at higher elevations. These projected climatic shifts align with the observed differences in ecosystem functioning and C dynamics across the scenarios. Our study, although focusing on only one growing season, found that the RCP 8.5 scenario led to marked differences in C fluxes and minor changes in leaf functional traits, mostly limited to a single species, of dominant alpine plants compared to both the RCP 4.5 and the Control scenarios. Contrary to our initial hypotheses, under moderate climate change (RCP 4.5), the ecosystem functioning remained consistent with the Control, showing no increase in its potential daytime C-sink strength. Under the more severe scenario (RCP 8.5), the data revealed a more complex response. Contrary to our hypothesis, no clear evidence of water limitation was observed, indicating that vegetation and soil processes remained relatively well sustained under the imposed conditions. The RCP 8.5 scenario strongly impacted physiological processes such as photosynthesis and respiration, with air temperature predominantly driving photosynthesis and soil temperature largely controlling respiration. Rising air temperature led to an average increase of 3.81 µmol mD^2^ sD^1^ in *GPP* and an increase of 2.49 µmol mD^2^ sD^1^ in *R*_*eco*_ compared to the Control, with no observed effect on *NEP* during the day. Plant photosynthesis in these C3 plants occurs only during daytime and is further limited to the growing season, when plants are exposed after snowmelt. In contrast, *R*_*eco*_ continues throughout the entire day and year. Rising daily minimum and maximum temperatures could therefore have different effects on ecosystem C uptake and release, with consequent influences on ecosystem C sequestration and feedback to climate change (Anderegg et al. 2015). Indeed, when considering the overall daily C balance, we found that nighttime *NEP*, driven solely by *R*_*eco*_, decreased under RCP 8.5 due to a pronounced increase in *R*_*eco*_. In contrast, during daytime, the effect of increased *R*_*eco*_ was compensated by high rates of *GPP*, resulting in similar *NEP* values across scenarios. Therefore, although RCP 8.5 conditions may initially enhance photosynthesis and result in a positive *NEP*, the overall daily C balance could become neutral or even negative over time, as the increased daytime C uptake was offset by enhanced nighttime *R*_*eco*_ under severe warmer conditions. However, soil basal respiration, measured at the same temperature for all treatments, did not increase in respect to Control, suggesting that, over this short period, soil microbial communities did not undergo thermal adaptation through shifts in community composition or physiological adjustments in response to warming. These observations are consistent with Alster et al. (2023), who highlighted that thermal adaptation of soil microbial respiration may require longer timescales or specific environmental conditions to become evident. To provide a broader and longer-term perspective, Wang et al. (2019) conducted a global meta-analysis of warming experiments across three grassland types—cold, temperate, and semi-arid—and reported distinct responses of C fluxes. In contrast to temperate and semi-arid grasslands, in cold grasslands, warming generally stimulated *GPP* more than *R*_*eco*_, leading to a net increase in C storage. However, this positive effect tended to weaken or disappear after more than three years of warming, likely due to resource limitations, shifts in plant and microbial communities, or reduced C use efficiency.

Taken together, these findings suggest that the balance between warming-induced increases in carbon uptake and *R*_*eco*_ is a key determinant of C cycling in alpine ecosystems and warrants further investigation over longer timescales.

### Plant–soil interactions and ecosystem functioning

Warming-induced changes in ecosystem functioning were accompanied by shifts in vegetation structure and carbon allocation patterns. Under RCP 8.5, we observed an increase in aboveground biomass accompanied by a decline in the belowground to aboveground biomass ratio under RCP 8.5. This pattern indicates a shift towards aboveground allocation, suggesting that plants may invest more resources into the development of leaves and stems to maximize photosynthetic capacity under conditions that favor aboveground growth. Similarly, Fazlioglu and Wan (2021) reported a significant increase in aboveground biomass of tundra and alpine plants with rising MAT, while total belowground and fine-root biomass either remained stable or declined across a broad MAT gradient under experimental warming (Björk et al. 2007; Gough and Hobbie, 2003). Such a shift implies that warming preferentially promotes shoot growth over root development, potentially modifying plant resource allocation strategies in cold ecosystems. Nevertheless, to overcome nutrient limitations when they occur, plants may increase root biomass to enhance soil exploration (Hodge 2004) or diversify the forms of N absorbed (Jacot et al. 2000).

Despite these structural changes, no detectable differences in soil nutrient concentrations were observed across treatments. This suggests that, over the short experimental period, nutrient availability did not differ significantly from the control, which may indicate that stronger biogeochemical feedbacks require longer timescales to emerge (Rustad et al. 2001; Melillo et al. 2017). Similar patterns have been reported in some warming studies (Sistla and Schimel, 2013; Bradford et al. 2008), where elevated temperatures enhanced C turnover without immediate changes in nutrient pools under short-term conditions.

Overall, early-stage warming appears to enhance plant productivity and carbon inputs to soil without immediate changes in nutrient pools, consistent with a transient decoupling between vegetation response and soil biogeochemistry.

### Species-specific responses and functional traits

The resilience of a plant community to climate change is determined by the individual sensitivity of species to environmental shifts, making it a species-specific process. Hence, predicting which species will benefit or decline under a changing climate is needed for accurate estimates of community composition changes and ecosystem functioning. We analyzed the responses of species, representative of different plant life forms, to various climate change scenarios, including forbs (*Potentilla aurea, Scorzoneroides montana*), graminoids (*Poa alpina, Carex foetida*), and dwarf shrub (*Salix herbacea*). Among them, only *Salix herbacea* showed detectable changes in leaf traits in response to climate change, particularly under the RCP 8.5 scenario. However, this did not coincide with an increase in its coverage, unlike the forbs *Potentilla aurea* and *Scorzoneroides montana*, which tended to expand, likely due to their more flexible functional traits that promote growth and spread under rising alpine temperatures (Steinbauer et al. 2022). This may indicate that *Salix herbacea* is undergoing functional adjustments that could enhance its competitiveness under future warming conditions, potentially preceding an expansion phase. Such a response is consistent with the broader concern that shrub encroachment may increasingly threaten alpine biomes, as expanding shrubs can reduce species richness through the replacement of herbaceous species (Elmendorf et al. 2012) or competitive exclusion (Boscutti et al. 2018).

*Salix herbacea* demonstrated a significant increase in its LMA and *N*_*a*_ and showed a tendency to increase in δ^1^DN. The ratio of leaf dry mass to leaf area reflects the cost of light interception at the leaf level; it is a key trait for understanding plant growth (Lambers and Poorter, 1992) and an important indicator of plant ecological strategies (Westoby et al. 2002). Indeed, species with a higher LMA, could exhibit longer leaf longevity due to increased leaf toughness, which enhances their ability to withstand extreme drought conditions (Li et al. 2021). Additionally, *Salix herbacea* exhibited the highest *N*_*a*_ levels, which may enhance its ability to conserve water while maintaining photosynthetic rates comparable to those of plants from wetter environments, as proposed by Wright et al. (2003). This phenomenon aligns with the *least-cost economic theory of photosynthesis* (Prentice et al. 2014, Wang et al. 2017), which suggests that plants optimize photosynthesis by balancing C assimilation efficiency with resource conservation. Plants may reduce stomatal conductance and transpiration rates under drought to minimize water loss while maintaining a steeper COD diffusion gradient, which sustains efficient C uptake into the leaves despite limited water availability (Flexas et al. 2004; Wang et al., 2017). The δ15N serves as a potential indicator of plant N metabolism and growth conditions (Serret et al., 2018). Increases in δ15N can signal altered soil N availability and changes in the N uptake pathway by plants, even when N concentrations remain unchanged. Generally, δ15N values rise with the opening of the ecosystem N cycle, defined as an increase in net N inputs and losses relative to internal N cycling (Hobbie and Högberg, 2012). For instance, in a subarctic bog, passive springtime warming using open-top chambers (soil temperature +1°C) led to higher δ15N values in *Andromeda polifolia* leaves, suggesting an increased reliance on less ^15^N-depleted inorganic N sources, possibly due to enhanced soil N mineralization rates (Aerts et al. 2009). However, no such change was observed with spring or summer warming for any other plant species in the same study, suggesting species-specific responses to climate manipulations.

### Conclusions

This study highlights how projected climate change scenarios may alter the structure and functioning of alpine grassland ecosystems. Simulating future conditions in controlled mesocosms—which can simultaneously manipulate multiple environmental parameters—provided insights into the complex and interactive mechanisms driving ecosystem functioning under projected climate change scenarios in alpine grassland.

Based on observations from a single growing season, our results indicate that alpine grassland ecosystems remain largely in a steady state under moderate warming scenarios (RCP 4.5). Contrary to our expectations, there were no significant changes in photosynthesis, *R*_*eco*_, nutrients concentrations or plant biomass under this scenario. In contrast, under the most severe climate scenario (RCP 8.5), the observed responses largely mirrored those hypothesized for the moderate warming scenario. Increases in *GPP* were supported by higher vegetation cover and plant biomass. This response was associated with a shift in plant biomass allocation strategies toward increased aboveground biomass and reduced belowground investment. Despite this enhanced C uptake, daytime *NEP* remained largely unchanged because increases in *GPP* were offset by a corresponding rise in *R*_*eco*_, with no detectable limitation of water and nutrients availability. However, because respiratory process operates continuously over the diel cycle, enhanced respiration under RCP 8.5 leads to increased C losses at night, potentially altering the balance between daytime C uptake and nighttime respiratory release on a diel basis.

Plant responses to climate change were species-specific, and among the dominant species, only *Salix herbacea* exhibited measurable adjustments of functional traits, indicating a greater capacity for physiological acclimation to climate change. These findings may help improve projections of ecosystem responses, thereby supporting the management and conservation of vulnerable ecosystems under climate warming.

## Acknowledgements

We are grateful to Luciano Spaccino and Fabio Trevisan for their assistance with stable isotope analyses and soil physicochemical characterization, respectively. We also thank Prof. Tanja Mimmo and Prof. Luigimaria Borruso for their scientific and financial support to the analytical work, data analysis, and Leonardo Latilla and all people involved in the monoliths transportation process.

This work was also made possible through the valuable collaboration and logistical support of Fondazione Montagna Sicura - Courmayeur, Valle d’Aosta, Aosta, Italy.

## Funding

This work was carried out within the framework of the Italian PRIN 2020 project MICROPLANTALP (*“MICROorganism–PLANT Interactions in the Forefield of Glaciers: a Hotspot for Studying the Impact of Climate Change in ALPine Habitats,”* 20204KF4RW). This study also benefited from the CNRS resources allocated to the French ECOTRONS Research Infrastructure, from the Occitanie Region and FEDER investments as well as from the state allocation ‘Investissement d’Avenir’ AnaEEFrance ANR-11-INBS-0001.

## Competing Interests

The authors declare that they have no known competing financial interests or personal relationships that could have appeared to influence the work reported in this paper.

## Notes

### Competing Interest Statement

The authors have declared no competing interest.

