## Supplementary material for "Severe warming alters biomass allocation and enhances ecosystem respiration in an alpine grassland: evidence from a short-term mesocosm experiment"

Federica D’Alò

Institute of Research on Terrestrial Ecosystems, National Research Council, Porano (TR), Italy

^†^ Author passed away before the submission of this manuscript

**Material and Methods**

**Soil physicochemical characterization**

Soil was collected at the beginning of the experiment and subsequently every 20 days using a sterilized spatula from at least three points within each mesocosm, up to a depth of 10 cm. Samples from the different points were mixed to obtain a composite sample and transferred into sterile Falcon tubes, which were immediately frozen and stored at −20 °C until analysis. To analyze the chemical composition of soil, samples were air-dried until they reached a constant weight and then sieved through a 2 mm mesh.

Soil pH was determined in both deionised water and CaCl₂ using a 1:2.5 soil-to-solution ratio. For each sample, 5 g of air-dried material was weighed into a clean, 50 mL centrifuge tube. To this, 12.5 mL of 0.01 M CaCl₂ or deionised water was added, and the mixture was shaken on a reciprocal shaker at room temperature for 15 minutes. The mixture was then centrifuged at 1000 rpm for 5 minutes. A calibrated pH meter (XS Instruments, Capri, Italy) was then immersed in the liquid phase, and once the meter stabilised, pH readings were recorded. An aliquot of the liquid phase from the deionised water extraction was transferred to a clean receptacle for nitrate (NO₃⁻) analysis using an ion chromatograph (Dionex ICS-6000 HPIC, Thermo Scientific, Germany). The IC system was equipped with a Dionex IonPac AS11-HC column (guard: 50 × 2 mm, 13 μm; analytical: 250 × 2 mm, 9 μm) and a conductivity detector with AERS 2 mm suppressor (29 mA, 0.38 mL min⁻¹). Eluent generation utilized an EGC 500 KOH cartridge (30 mM, isocratic), while the injection volume was 10 μL.

To measure ammonium (NH₄⁺), 1 mL of the extract in CaCl₂ was transferred into a 15 mL Falcon tube and analysed following the Berthelot method with sodium salicylate (Kempers and Zweers, 1986). Briefly, 2.50 mL of Berthelot reagent (A) was added to the extract, and the mixture was shaken intermittently for 15 minutes. Subsequently, 2.50 mL of Berthelot reagent (B) was added to all Falcon tubes, which were then sealed and left to stand for 30 minutes. 1 mL of the resulting mixture was transferred into cuvettes and measured with an Agilent Cary Series UV-Vis spectrophotometer (Agilent Technologies, Santa Clara, CA, USA) at a wavelength of 660 nm.

Soil available phosphate (PO₄³⁻) was determined following the Olsen method (Murphy and Riley, 1962). 1 g of soil with a small amount of activated charcoal was mixed in a 50 mL centrifuge tube, along with 20 mL of 0.5 M NaHCO₃. A 5.0 mL aliquot of the filtrate was transferred to a 50 mL volumetric flask, acidified with 2 drops of 0.25% p-nitrophenol and 2.5 M H₂SO₄ until the yellow color disappeared, and gently shaken for approximately 10 minutes to expel CO₂. 8 mL of Murphy and Riley reagent was added to each flask, which was then filled to 50 mL with distilled water and allowed to stand for 10 minutes. 1 mL of the resulting mixture was transferred into cuvettes and measured with an Agilent Cary Series UV-Vis spectrophotometer (Agilent Technologies, Santa Clara, CA, USA) at a wavelength of 720 nm.

For total C (TC), total nitrogen (TN), and elemental composition, a subsample of soil was homogenized in a Mixer Mill MM 400 (Retsch, Germany). To determine TOC and TN, soil samples were analysed using the Flash EA 112 Elemental Analyser (Thermo Scientific, Germany).

For element analysis, 1 g of fine, homogeneous soil was placed into a polyethylene container with both the top and bottom covered by polypropylene film specifically designed for XRF analysis. The samples were analysed using an X-ray fluorescence spectrometer (Epsilon 4 XRF Spectrometer, Malvern Panalytical, Netherlands) with a standard acquisition time (20 min). For a representative subset of samples, loss on ignition (LOI) was determined by heating 1 g of soil for 4 hours at 900°C. The LOI values were then incorporated into the processing software to enhance the accuracy of elemental quantification.

**Supplementary tables**

**Table 1S** Results of the nonlinear mixed-effects model evaluating the effects of climate scenario, sampling date, their interaction, and pre-treatment values (included to account for initial variability) on soil physicochemical parameters. TC, total carbon (%); TN, total nitrogen (%), NH_4_^+^, ammonium ion (mg/kg); NO_3_^-^, nitrate ion (mg/kg); TP, total phosphorus (%); PO₄³⁻, phosphate ion (mg/kg). The table reports degrees of freedom (numDF and denDF), F-values, and p-values for fixed effects and their interactions.

|  | numDF | denDF | F-value | p-value |
| --- | --- | --- | --- | --- |
| pH |  |  |  |  |
| Intercept | 1 | 18 | 3728.363 | <0.0001 |
| Climate scenario | 2 | 6 | 2.057 | 0.2088 |
| Date | 2 | 18 | 83.236 | <0.0001 |
| Pre-treatment pH | 1 | 6 | 0.560 | 0.4827 |
| Climate scenario x date | 4 | 18 | 6.994 | 0.0014 |
| TC |  |  |  |  |
| Intercept | 1 | 18 | 640.7138 | <0.0001 |
| Climate scenario | 2 | 6 | 0.5029 | 0.6282 |
| Date | 2 | 18 | 8.4947 | 0.0025 |
| Pre-treatment TC | 1 | 6 | 28.9227 | 0.0017 |
| Climate scenario x date | 4 | 18 | 1.7497 | 0.1831 |
| TN |  |  |  |  |
| Intercept | 1 | 18 | 1361.3382 | <0.0001 |
| Climate scenario | 2 | 6 | 0.7861 | 0.4975 |
| Date | 2 | 18 | 8.1592 | 0.0030 |
| Pre-treatment TN | 1 | 6 | 20.0146 | 0.0042 |
| Climate scenario x date | 4 | 18 | 2.1068 | 0.1221 |
| C:N ratio |  |  |  |  |
| Intercept | 1 | 18 | 6536.528 | <0.0001 |
| Climate scenario | 2 | 6 | 2.512 | 0.1613 |
| Date | 2 | 18 | 2.350 | 0.1239 |
| Pre-treatment C:N ratio | 1 | 6 | 0.872 | 0.3865 |
| Climate scenario x date | 4 | 18 | 2.305 | 0.0979 |
| NH_4_^+^ |  |  |  |  |
| Intercept | 1 | 18 | 387.6125 | <0.0001 |
| Climate scenario | 2 | 6 | 0.9140 | 0.4503 |
| Date | 2 | 18 | 18.6196 | <0.0001 |
| Pre-treatment NH_4_^+^ | 1 | 6 | 13.3623 | 0.0106 |
| Climate scenario x date | 4 | 18 | 1.3137 | 0.3026 |
| NO_3_^-^ |  |  |  |  |
| Intercept | 1 | 18 | 1362.5681 | <0.0001 |
| Climate scenario | 2 | 6 | 3.5044 | 0.0981 |
| Date | 2 | 18 | 2.9598 | 0.0774 |
| Pre-treatment NO_3_^-^ | 1 | 6 | 9.0715 | 0.0236 |
| Climate scenario x date | 4 | 18 | 2.5572 | 0.0743 |
| TP |  |  |  |  |
| Intercept | 1 | 18 | 214.6216 | <0.0001 |
| Climate scenario | 2 | 6 | 1.5622 | 0.2843 |
| Date | 2 | 18 | 0.8178 | 0.4572 |
| Pre-treatment TP | 1 | 6 | 8.2238 | 0.0285 |
| Climate scenario x date | 4 | 18 | 1.0958 | 0.3885 |
| PO_4_^3-^ |  |  |  |  |
| Intercept | 1 | 18 | 449.2865 | <0.0001 |
| Climate scenario | 2 | 6 | 17.7862 | 0.0030 |
| Date | 2 | 18 | 2.6861 | 0.0953 |
| Pre-treatment PO_4_^3-^ | 1 | 6 | 13.5809 | 0.0103 |
| Climate scenario x date | 4 | 18 | 1.1739 | 0.3554 |

_TC, total carbon (%); TN, total nitrogen (%), NH4+, ammonium ion (mg/kg); NO3-, nitrate ion (mg/kg); TP, total phosphorus (%); PO₄³⁻, phosphate ion (mg/kg). The table reports degrees of freedom (numDF and denDF), F-values, and p-values for fixed effects and their interactions._

**Table 2S** Results of a nonlinear mixed-effects model used to analyze C fluxes and vegetation coverage as a function of climate scenario and date of sampling variables.

|  | numDF | denDF | F-value | p-value |
| --- | --- | --- | --- | --- |
| R_eco_ |  |  |  |  |
| Intercept | 1 | 90 | 35.66109 | <0.0001 |
| Climate scenario | 2 | 7 | 11.68740 | 0.0059 |
| Date | 10 | 90 | 17.28331 | <0.0001 |
| Climate scenario x date | 20 | 90 | 4.18163 | <0.0001 |
| R_eco_ | **Fitted coefficients** | **Std.Error** | **denDF** | **p-value** |
| Intercept | 2.2014828 | 0.7386386 | 90 | 0.0037 |
| RCP 4.5 | 0.8940217 | 0.5589099 | 7 | 0.1537 |
| RCP 8.5 | 2.4915300 | 0.5589099 | 7 | 0.0029 |
| GPP | **numDF** | **denDF** | **F-value** | **p-value** |
| Intercept | 1 | 90 | 31.63361 | < 0.0001 |
| Climate scenario | 2 | 7 | 2.05105 | 0.1990 |
| Date | 10 | 90 | 10.61335 | < 0.0001 |
| Climate scenario x date | 20 | 90 | 2.49530 | 0.0018 |
| GPP | **Fitted coefficients** | **Std.Error** | **denDF** | **p-value** |
| Intercept | 10.650024 | 2.3990365 | 90 | 0.0000 |
| RCP 4.5 | 0.704988 | 1.7806682 | 7 | 0.7040 |
| RCP 8.5 | 3.819940 | 1.7806682 | 7 | 0.0691 |
| NEP | **numDF** | **denDF** | **F-value** | **p-value** |
| Intercept | 1 | 90 | 31.400043 | <0.0001 |
| Climate scenario | 2 | 7 | 0.044689 | 0.9566 |
| Date | 10 | 90 | 6.344069 | <0.0001 |
| Climate scenario x date | 20 | 90 | 2.199006 | 0.0063 |
| NEP | **Fitted coefficients** | **Std.Error** | **denDF** | **p-value** |
| Intercept | 8.453 | 1.686 | 90 | 0.000 |
| RCP 4.5 | -0.298 | 1.607 | 7 | 0.858 |
| RCP 8.5 | 1.424 | 1.697 | 7 | 0.405 |
| Vegetation coverage | **numDF** | **denDF** | **F-value** | **p-value** |
| Intercept | 1 | 54 | 200.981 | <0.0001 |
| Climate scenario | 2 | 7 | 2.993 | 0.126 |
| Date | 6 | 54 | 4.758 | 0.001 |
| Pre-treatment vegetation coverage | 1 | 6 | 0.282 | 0.615 |
| Climate scenario x date | 12 | 54 | 0.525 | 0.889 |
| Vegetation coverage | **Fitted coefficients** | **Std.Error** | **denDF** | **p-value** |
| Intercept | 60.675 | 23.525 | 54 | 0.013 |
| RCP 4.5 | 0.906 | 14.263 | 6 | 0.951 |
| RCP 8.5 | 28.621 | 13.886 | 6 | 0.085 |

_For vegetation coverage, the model also included the pre-treatment differences as a covariate to account for initial variability before the application of climate scenarios. The table reports degrees of freedom (numDF and denDF), F-values, and p-values for fixed effects and their interactions. Estimated fixed effect coefficients for each climate scenario (relative to the control) are shown with standard errors (Std. Error), degrees of freedom (denDF), and p-values._

**Table 3S** Summary of model-averaging results showing the relative importance of environmental predictors for C fluxes.

|  | Intercept | T soil | DF | logLik | AIC | ΔAIC | AIC weight |
| --- | --- | --- | --- | --- | --- | --- | --- |
| R_eco_ | 0.028 | 0.181 | 10 | -127.139 | 276.1 | 0 | 1 |
| GPP | 9.961 | 0.133 | 10 | -245.889 | 513.6 | 2.99 | 0.183 |
| NEP | 9.747 | 0.045 | 10 | -221.793 | 465.4 | 2.58 | 0.216 |
|  | **Intercept** | **T air** | **DF** | **logLik** | **AIC** | **ΔAIC** | **AIC weight** |
| GPP | 10.330 | 0.149 | 10 | -245.567 | 513.0 | 2.34 | 0.236 |
| NEP | 9.422 | -0.027 | 10 | -219.554 | 460.9 | 0 | 0.721 |

_Only models with ΔAIC ≤ 4 are considered. The table reports the intercept, coefficients for soil and air temperature (T soil and T air), degrees of freedom (DF), log-likelihood (logLik), Akaike Information Criterion (AIC), AIC difference (ΔAIC), and AIC weight (AIC weight) for each model. The best predictor was identified based on the model with the lowest ΔAIC (≤ 4) and the highest AIC weight, indicating the model with the strongest support_.

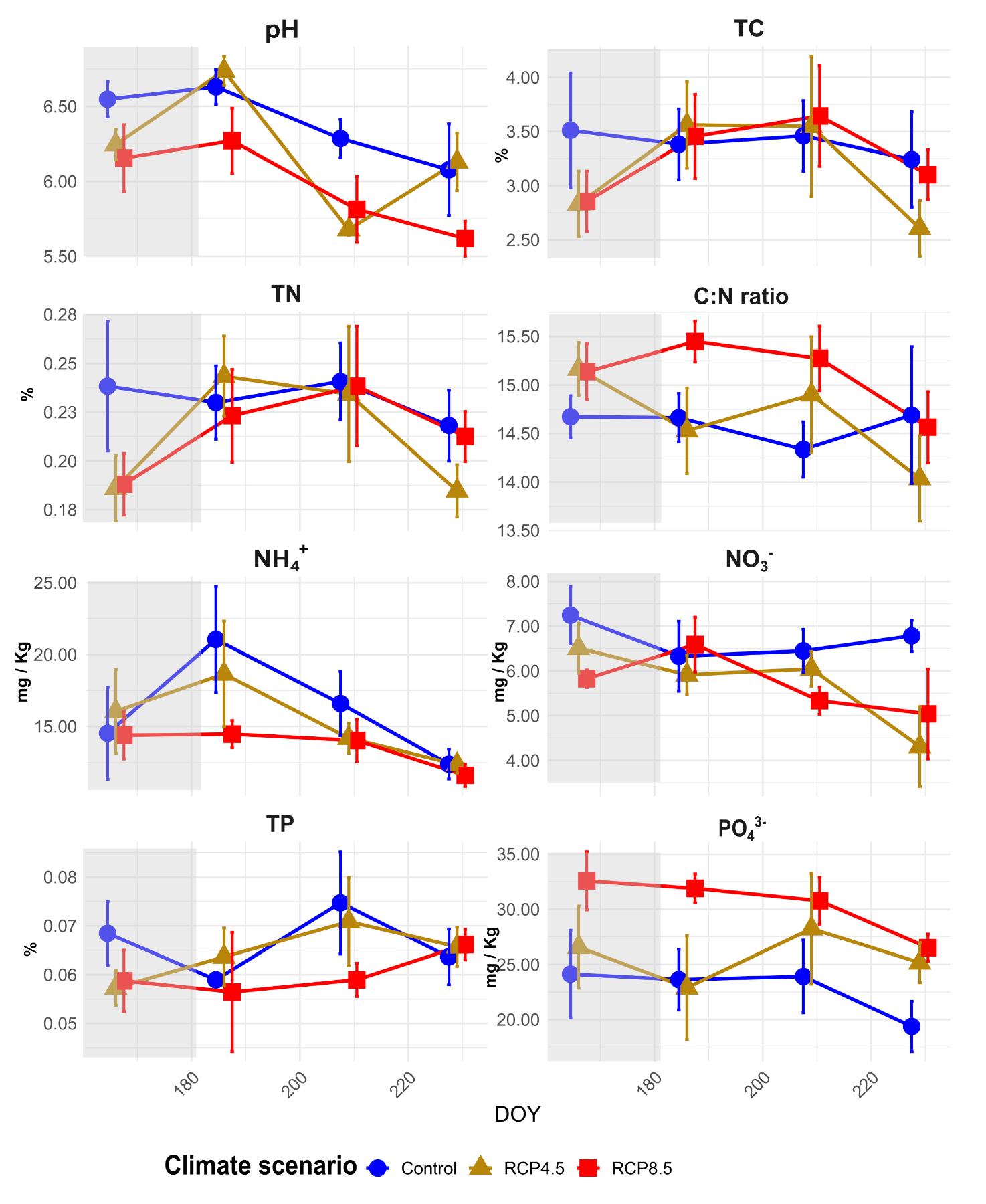

**Supplementary Fig 1** **Trends in means ± standard error (n = 4) of edaphic parameters under different climate scenarios throughout the short-term experiment.** TC, total carbon (%); TN, total nitrogen (%), NH_4_^+^, ammonium ion (mg/Kg); NO_3_^-^, nitrate ion (mg/Kg); TP, total phosphorus (%); PO₄³⁻, phosphate ion (mg/Kg). The gray section represents the two-week period prior to the start of measurements. DOY = Day Of Year
